# Colonization of the tsetse fly midgut with commensal *Enterobacter* inhibits trypanosome infection establishment

**DOI:** 10.1101/472373

**Authors:** Brian L. Weiss, Michele A. Maltz, Aurélien Vigneron, Yineng Wu, Katharine Walter, Michelle B. O’Neill, Jingwen Wang, Serap Aksoy

## Abstract

Tsetse flies (*Glossina* spp.) vector pathogenic trypanosomes (*Trypanosoma* spp.) in sub-Saharan Africa. These parasites cause human and animal African trypanosomiases, which are debilitating diseases that inflict an enormous socio-economic burden on inhabitants of endemic regions. Current disease control strategies rely primarily on treating infected animals and reducing tsetse population densities. However, relevant programs are costly, labor intensive and difficult to sustain. As such, novel strategies aimed at reducing tsetse vector competence require development. Herein we investigated whether an *Enterobacter* bacterium (*Esp_Z*), which confers *Anopheles gambiae* with resistance to *Plasmodium*, is able to colonize tsetse and induce a trypanosome refractory phenotype in the fly. *Esp_Z* established stable infections in tsetse’s gut, and exhibited no adverse effect on the survival of individuals from either group. Flies with established *Esp_Z* infections in their gut were significantly more refractory to infection with two distinct trypanosome species (*T. congolense*, 6% infection; *T. brucei*, 32% infection) than were age-matched flies that did not house the exogenous bacterium (*T. congolense*, 36% infected; *T. brucei*, 70% infected). Additionally, 52% of *Esp_Z* colonized tsetse survived infection with entomopathogenic *Serratia* marcescens, compared with only 9% of their wild-type counterparts. These parasite and pathogen refractory phenotypes result from the fact that *Esp_Z* acidifies tsetse’s midgut environment, which inhibits trypanosome and *Serratia* growth and thus infection establishment. Finally, we determined that *Esp_Z* infection does not impact the fecundity of male or female tsetse, nor the ability of male flies to compete with their wild-type counterparts for mates. We propose that *Esp_Z* could be used as one component of an integrated strategy aimed at reducing the ability of tsetse to transmit pathogenic trypanosomes.

**Author Summary:** Tsetse flies transmit pathogenic African trypanosomes, which are the causative agents of socio-economically devastating human and animal African trypanosomiases. These diseases are currently controlled in large part by reducing the population size of tsetse vectors through the use of insecticides, traps and sterile insect technique. However, logistic and monetary hurdles often preclude the prolonged application of procedures necessary to maintain these control programs. Thus, novel strategies, including those aimed at sustainably reducing the ability of tsetse to transmit trypanosomes, are presently under development. Herein we stably colonize tsetse flies with a bacterium (*Enterobacter* sp. Z, *Esp_Z*) that acidifies their midgut, thus rendering the environment inhospitable to infection with two distinct, epidemiologically important trypanosome strains as well as an entomopathogenic bacteria. In addition to inducing a trypanosome refractory phenotype, colonization of tsetse with *Esp_Z* exerts only a modest fitness cost on the fly. Taken together, these findings suggest that *Esp_Z* could be applied to enhance the effectiveness of currently employed tsetse control programs.

## Introduction

Insects transmit numerous vertebrate pathogens that cause devastating disease throughout tropical and subtropical regions around the globe. The lack of effective and affordable vaccines, coupled with insect and pathogen resistance to pesticides and drug treatments, respectively, severely limits disease control. Many vertebrate pathogens are acquired by insect vectors via the ingestion of an infectious blood meal. The disease causing agent must then establish an infection in the insect’s gut prior to being transmitted to a new vertebrate host during a subsequent bite. In most cases pathogens are eliminated from the insect vector prior to transmission to a new vertebrate host. This outcome reflects the presence of dynamic active and passive immune barriers that function locally in the insect gut and systemically in the hemocoel (Baxter et al., 2017; Saraiva et al., 2016; Aksoy et al., 2013).

Although few insect vectors support transmissible infections with vertebrate pathogens, all house symbiotic microorganisms in their gut that influence numerous aspects of their host’s physiological homeostasis. Symbiotic associations between arthropod disease vectors and enteric bacteria have been particularly well-studied in an effort to determine how these microbes influence their host’s ability to transmit disease (Weiss and Aksoy, 2011; Cirimotich et al., 2011; Narasimhan and Fikrig, 2015; Song et al., 2018; Dey et al., 2018). Tsetse flies, which are the prominent vectors of pathogenic African trypanosomes, house a taxonomically diverse enteric microbiota that includes endosymbiotic *Wigglesworthia* and *Sodalis* (Wang et al., 2013) as well as an assemblage of bacteria obtained from the fly’s environment (Aksoy et al., 2014; Geiger et al., 2011; Lindh and Lehane, 2011). Both *Wigglesworthia* and *Sodalis* are maternally transmitted to developing intrauterine larvae during tsetse’s unique mode of viviparous reproduction (Wang et al. 2013; Benoit et al., 2015). *Wigglesworthia* influences trypanosome infection establishment in tsetse by regulating the production of trypanocidal PGRP-LB (Wang et al., 2009; Wang et al., 2012). Additionally, tsetse that undergo larval development in the absence of this bacterium fail to synthesize a gut-associated peritrophic matrix during adulthood (Weiss et al., 2013) This structure is an important mediator of tsetse’s vector competence because it serves as a physical barrier that ingested parasites must traverse in order to successfully colonize the fly’s gut (Weiss et al., 2014) and subsequently the salivary glands for transmission in saliva (Vigneron et al., 2018). *Sodalis’* impact on tsetse vector competency is less known, although studies suggest that a positive correlation exists between the prevalence and density of this bacterium and trypanosome infection prevalence (Welburn et al., 1993; Dale and Welburn, 2001; Farikou et al., 2010; Soumana et al., 2013; Griffith et al., 2018). Like mosquitoes, tsetse’s gut also harbors a diverse population of bacteria obtained from the fly’s environment (Geiger et al., 2011; Lindh and Lehane, 2011; Aksoy et al., 2014). However, the effect of these bacteria on tsetse vector competency is poorly understood.

Mosquitoes, including *Anopheles gambiae* and *Aedes aegypti*, also house bacteria in their gut, and these microbes play a significant role in the ability of their host to transmit vertebrate pathogens. Boissiere et al. (2012) discovered a positive correlation between the density of enteric Enterobacteriaceae and *Plasmodium* infection prevalence in field-captured *An. gambiae*. These midgut microbes, as well as the enteric microbiota found in *Ae. aegypti*, indirectly regulate their host’s vector competency by modulating basal expression of genes that encode anti-*Plasmodium* and anti-dengue effector proteins (Cirimotich et al., 2011; Dennison et al., 2014; Bahia et al., 2014). Other members of the mosquito enteric microbiota exert direct effects on their host’s vector competency. Specifically, a *Chromobacterium* isolated from *Ae. aegypti* secretes factors that exhibit anti-*Plasmodium* and anti-Dengue activity (Ramirez et al., 2014). Also, laboratory reared *A. gambiae* present an abnormal *Plasmodium* refractory phenotype when their guts are colonized with a strain of *Enterobacter* (*Esp_Z*) that had been previously isolated from field-captured mosquitoes. *Esp_Z* was determined to produce reactive oxygen intermediates (ROIs) that exhibit direct anti-*Plasmodium* properties (Cirimotich et al., 2011; Dennison et al., 2016).

In this study we investigated whether *Esp_Z* isolated from *A. gambiae* is able to successfully colonize tsetse’s gut and induce parasite and pathogen refractory phenotypes in the fly. We found that this bacterium can reside stably in tsetse’s midgut without imparting a detrimental fitness cost on the fly. *Esp_Z* colonized tsetse present an acidified midgut environment that is inhospitable to both African trypanosomes and entomopathogenic *Serratia marcescens*. We discuss the potential utility of *Esp_Z* as a novel component of currently used area wide integrated pest management strategies aimed at controlling tsetse populations and thus transmission of African trypanosomes.

## Results

### Bacterial infection outcomes in tsetse’s midgut, and subsequent fly survival

We investigated the ability of *Esp_Z* and *Sodalis* (as a control) to colonize the gut of both wild-type (hereafter referred to as ‘*Gmm*^WT^’) and symbiont-free tsetse (aposymbiotic, hereafter referred to as ‘*Gmm*^Apo^’). *Gmm*^WT^ flies were used to interrogate the interaction between *Esp_Z* and the natural tsetse microbiota, while the use of *Gmm*^Apo^ individuals allowed us to correlate the presence of distinct, experimentally introduced bacterial taxa with specific fly phenotypes. We challenged all flies *per os* with 1×10^3^ CFU of either *Esp_Z* or *Sodalis* in their first blood meal and then monitored bacterial proliferation over an 28 day period. By 7 days post-inoculation, midgut bacterial density in *Gmm*^WT^ that housed *Esp_Z* (*Gmm*^WT/*Esp_Z*^) and *Sodalis* (*Gmm*^WT/*Sgm*^) was 1.9×10^7^ ± 6.4×10^6^ CFU and 4.5×10^5^ ± 6.4×10^6^ CFU, respectively, and *Gmm*^Apo/*Esp_Z*^ (9.3×10^6^ ± 5.3×10^5^ CFU) and *Gmm*^Apo/*Sgm*^ (1.4×10^6^ ± 3.9×10^5^ CFU) flies harbored a similar bacterial density at the same time point post-inoculation (Fig. 1A). The midgut density of *Esp_Z* and *Sodalis* did not change significantly in any of the fly groups over the following 21 days (Fig. 1A), thus suggesting that the bacteria had achieved stable-state infections within their fly hosts by one week post-acquisition.

**Figure 1.**
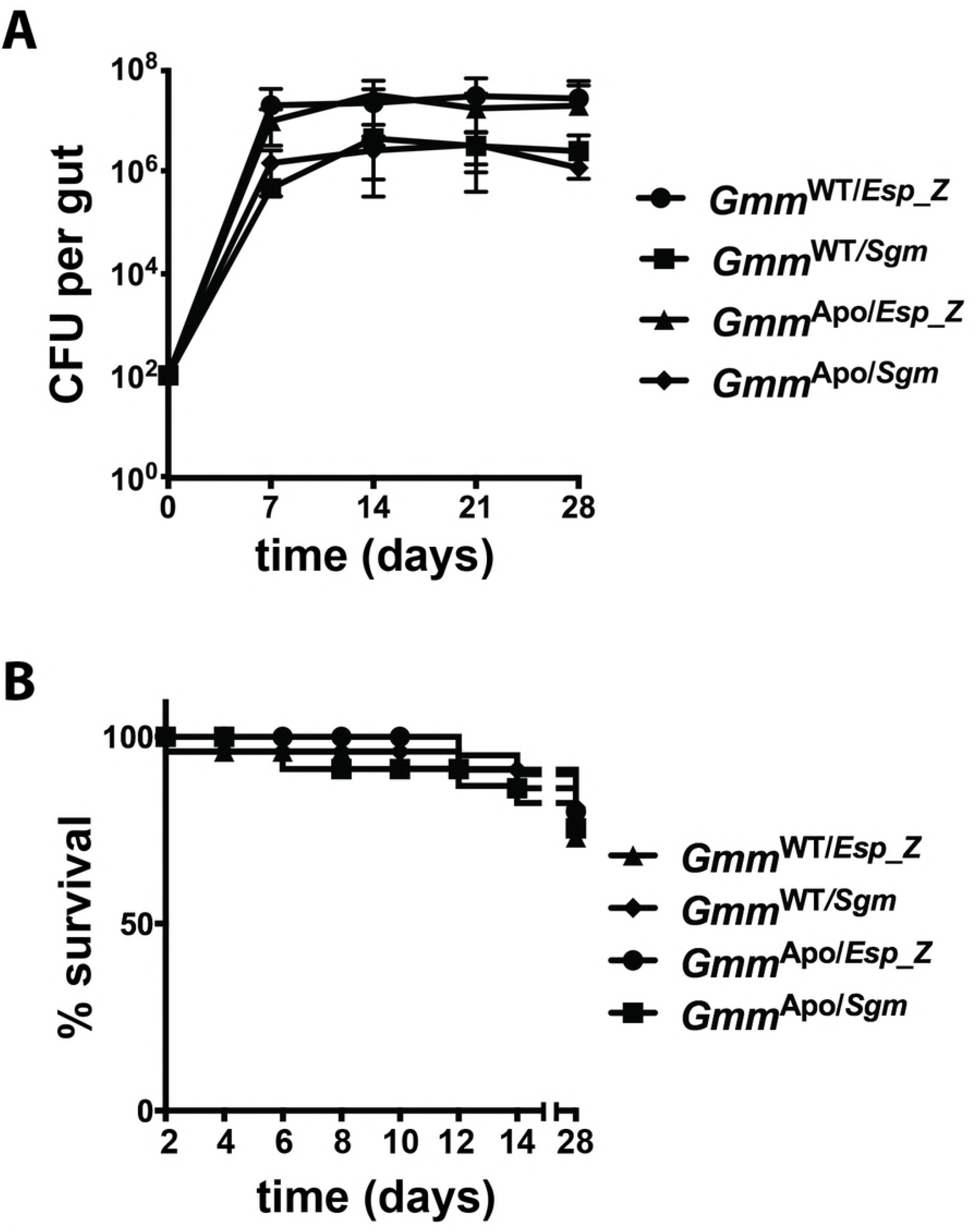
Bacterial colonization of tsetse’s midgut, and the effect on fly survival. Distinct groups of newly emerged adult wild-type (*Gmm*^WT^) and aposymbiotic (*Gmm*^Apo^) females were colonized with 1×10^3^ CFU of either *Enterobacter* (*Esp_Z*) or *Sodalis* (*Sgm*), and then bacterial density and fly survival was measured. (A) Average number (±SEM) of bacterial CFUs per tsetse gut per group per time point. *n*≥5 individuals per group per timepoint. (B) Kaplan-Meier plot depicting survival of *Gmm*^WT^ and *Gmm*^Apo^ females colonized with either *Esp_Z* or *Sgm*. Infection experiments were performed in triplicate, using 25 flies per replicate. No significant difference in survival was observed between any of the fly groups (*p*=0.88; log-rank test).

We next examined whether midgut infections with *Esp_Z* or *Sodalis* impacted tsetse survival. We found that 76% of *Gmm*^WT/*Esp_Z*^ and 84% of *Gmm*^WT/*Sgm*^ survived for 28 days following bacterial inoculation. Similarly, 84% and 80% of *Gmm*^Apo/*Esp_Z*^ and *Gmm*^Apo/*Sgm*^, respectively, survived the duration of the experiment (Fig. 1B). Percent survival was not significantly different between any of these groups, indicating that *Esp_Z* and *Sodalis* both exhibit commensal phenotypes in wild-type and aposymbiotic tsetse.

### Enterobacter is resistant to Peptidoglycan Recognition Protein-LB (PGRP-LB)

The midgut of adult tsetse expresses *peptidoglycan recognition protein LB* (*pgrp-lb*), which encodes a pattern recognition receptor that exhibits potent antimicrobial activity (Wang et al., 2009; Wang et al., 2012). Thus, in order to colonize tsetse’s midgut, a microorganism must be resistant to this molecule. We investigated whether innate resistance to PGRP-LB represents one mechanism that allows *Esp_Z* to colonize tsetse’s gut. We found that 108% (±16) of *Esp_Z* cells were able to survive 1 h in the presence of recPGRP-LB, while only 2.3% (±1.0) *E. coli* cells survived for the same time period. Additionally, 154% (±14) of *Sodalis* cells survived following a 12 h incubation with recPGRP-LB (Fig. 2). These findings suggest that like native *Sodalis, Esp_Z* is resistant to the antimicrobial properties of PGRP-LB and is able to replicate in the presence of this protein (as indicated by an increase in bacterial density compared to the initial inoculate). This phenotype may facilitate this bacterium’s ability to successfully colonize tsetse’s gut.

**Figure 2.**
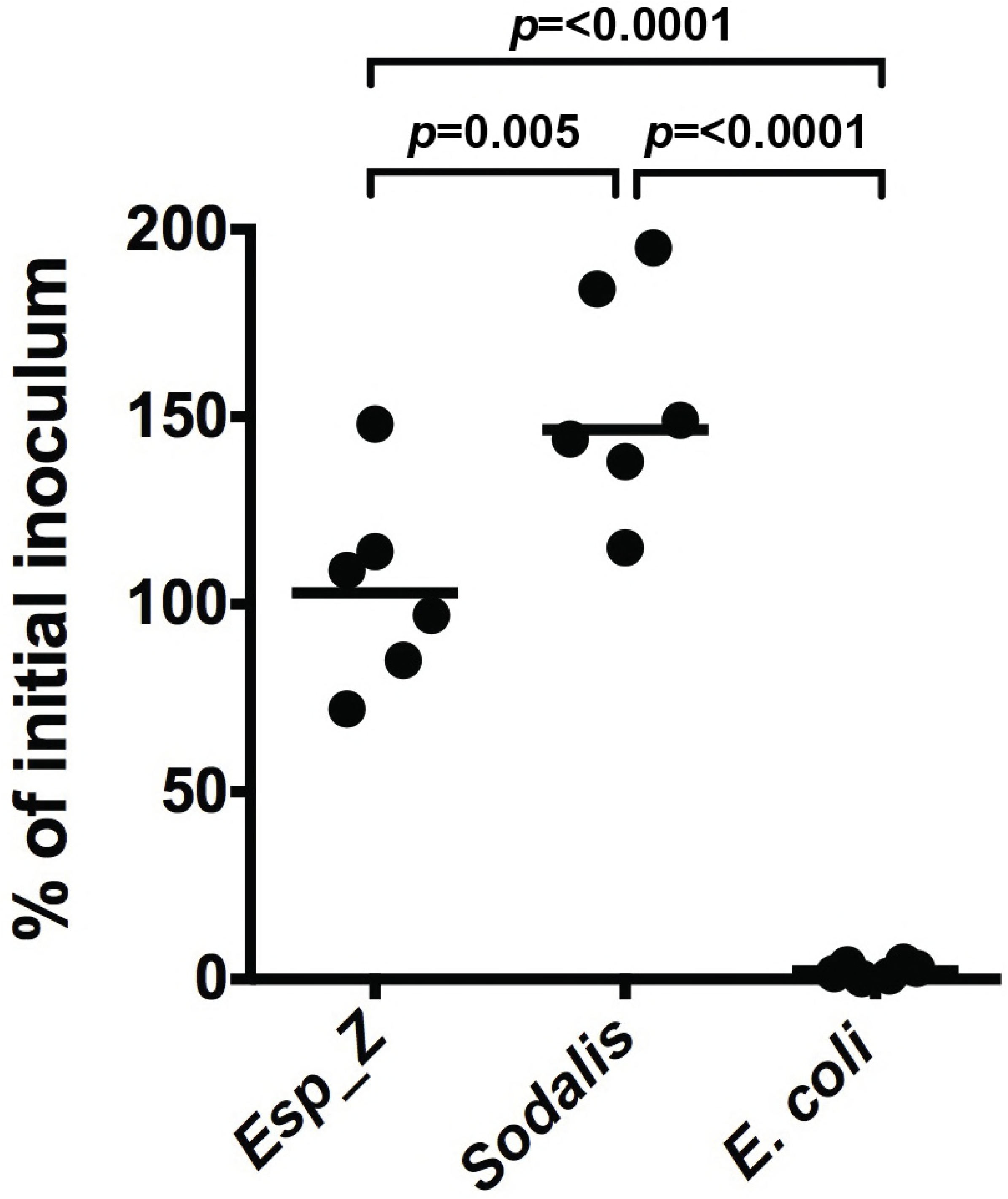
*Esp_Z* is resistant to normally bactericidal Peptidoglycan Recognition Protein-LB. Survival of cultured *Esp_Z, Sodalis* and *E. coli* following exposure (1 hr for *Esp_Z* and *E. coli*, and 24 hr for *Sodalis*) to recombinant (rec) PGRP-LB (10 μg/ml). Results are presented as % of initial inoculum, which was determined by dividing the number of bacterial CFU present after treatment with recPGRP-LB by the number of CFU present prior to inoculation. Statistical significance was determined using a one-way ANOVA followed by Tukey’s HSD post-hoc analysis

### Esp_Z colonized aposymbiotic tsetse present a trypanosome refractory phenotype

*Esp_Z* successfully colonizes the gut of *Gmm*^Apo^ flies, resides in the niche for at least 28 days, and has no impact on fly survival during that time period. Thus, we next evaluated whether colonization with this bacterium impacts trypanosome infection establishment in tsetse’s midgut. We began by challenging mature *Gmm*^Apo^ because they are highly susceptible to trypanosome infection (~50%) while their age-matched *Gmm*^WT^ counterparts are refractory (~3%) (Weiss et al., 2013). Distinct groups of eight day old *Gmm*^Apo/*Sgm*^ and *Gmm*^Apo/*Esp_Z*^, which housed similar numbers of their respective exogenous bacteria (S1 Fig. A), were administered a meal supplemented with 1×10^6^ blood stream form (BSF) trypanosomes per ml of blood. Thereafter all flies were maintained on regular blood for two weeks, at which point their midguts were dissected and microscopically examined for the presence of parasites. An age-matched control cohort consisted of similarly challenged *Gmm*^Apo^ flies. We found that infection prevalence in the *Gmm*^Apo/*Sgm*^ group (57%) was similar to that of *Gmm*^Apo^ controls (52%), while infection prevalence in *Gmm*^Apo/*Esp_Z*^ individuals was significantly lower (19%) (Fig. 3A). These data indicate that the presence of *Esp_Z* in tsetse’s gut interferes with the ability of trypanosomes to establish an infection in this niche. This parasite resistant phenotype is similar to that which occurs in the gut of *Esp_Z* colonized mosquitoes following exposure to malaria parasites (Cirimotich et al., 2011; Dennison et al., 2016).

**Figure 3.**
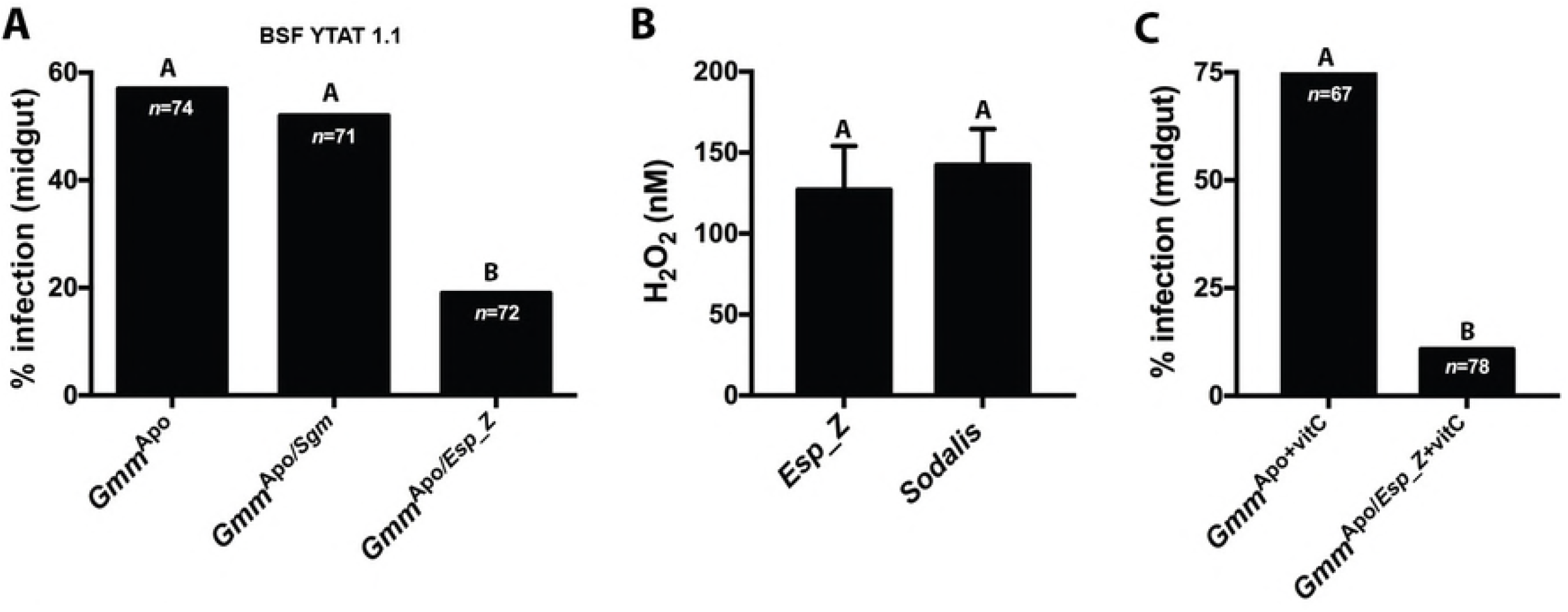
The trypanosome refractory phenotype exhibited by *Gmm*^Apo/*Esp_Z*^ flies is not directly caused by reactive oxygen intermediates. (A) Percentage of *Gmm*^Apo^, *Gmm*^Apo/*Sgm*^ and *Gmm*^Apo/*Esp_Z*^ flies harboring midgut infections with bloodstream form (BSF) YTAT 1.1 trypanosomes. Statistical analysis was performed using a GLM followed by multiple comparisons and Tukey contrasts, and different letters represents statistical significance between treatments and controls. (B) Mid-log phase cultures of *Esp_Z* and *Sodalis* synthesize similar quantities of H_2_O_2_ (*p*=0.6; paired t-test). Measurements were taken from 7 distinct clonal populations of each bacterium. (C) Percentage of *Gmm*^Apo^ and *Gmm*^Apo/*Esp_Z*^ flies infected with BSF YTAT 1.1 trypanosomes after being fed blood meals containing antioxidant vitamin C over the course of the 14 day experiment. Despite exposure of both tsetse groups to the ROI-suppressing vitamin, *Gmm*^Apo/*Esp_Z*^ flies were still significantly more refractory to trypanosome infection than were *Gmm*^Apo^ individuals (*p*=0.002; GLM Wald test). In (A), (B) and (C) different letters represents statistical significance between treatments and treatments and controls.

### African trypanosomes are not susceptible to Esp_Z generated reactive oxygen intermediates

In *A. gambiae, Esp_Z* produces reactive oxygen intermediates (ROIs) that are directly toxic to *Plasmodium* (Cirimotich et al., 2011; Dennison et al., 2016). ROIs have also been implicated as mediators of trypanosome infection outcomes in tsetse. Specifically, tsetse are rendered susceptible to trypanosome infection when the initial infectious blood meal is supplemented with the antioxidants vitamin C or cysteine (MacLeod et al., 2007; Vigneron et al., 2018). These antioxidants detoxify ROIs that otherwise induce programmed cell death processes in trypanosomes (Ridgely et al., 1999). In light of this information, we investigated the correlation between *Esp_Z* generated ROIs and the trypanosome refractory phenotypes we observed in adult *Gmm*^Apo/*Esp_Z*^. As an indicator of bacterial ROI production, we quantified H_2_O_2_ concentrations in supernatants from mid-log phase *Esp_Z* and *Sodalis* cultures. *Esp_Z* and *Sodalis* supernatants contained 127nM (±15) and 142nM (±13) of H_2_O_2_, respectively (Fig. 3B).

We next tested whether ROIs produced by *Esp_Z* inhibit the ability of trypanosome to infect *Gmm*^Apo/*Esp_Z*^. Individual groups of eight day old *Gmm*^Apo^ and *Gmm*^Apo/*Esp_Z*^ were offered a blood meal containing infectious trypanosomes together with the antioxidant vitamin C. All trypanosome challenged flies were subsequently maintained on vitamin C supplemented blood for 14 days. Under these conditions, 74% of *Gmm*^Apo+vitC^ were infected with trypanosomes, while only 11% of their *Gmm*^Apo/*Esp_Z*+vitC^ counterparts housed parasite infections (Fig. 3C). These results suggest that ROIs produced by *Esp_Z* that reside stably in tsetse’s gut are not the sole determinants of the fly’s susceptibility to infection with trypanosomes.

### Enterobacter produces acid that is toxic to trypanosomes

We observed that *Gmm*^Apo/*Esp_Z*^ are significantly more refractory to infection with trypanosomes than are *Gmm*^Apo/*Sgm*^ individuals, despite the fact that *Sodalis* and *Esp_Z* produce similar amounts of H_2_O_2_. This outcome implies that *Esp_Z* modulates trypanosome infection outcomes in tsetse via a mechanism other than ROI production. Several enteric commensals, including *Enterobacter* spp. (Podlesny et al., 2017; Fischer et al., 2017; Goldford et al., 2018), produce organic acids, and these products can inhibit pathogen growth by creating an acidic environment (Buffie et al., 2013; Neal-McKinney et al., 2012). Because many trypanosomatids, including members of the genera *Trypanosoma* and *Leishmania*, are highly sensitive to environmental pH (Nolan et al., 2000; Zilberstein and Shapira, 1994), we investigated whether *Esp_Z* creates an acidic environment that prohibits *T. brucei* growth *in vitro* and *in vivo*. Specifically, we added heat killed (HK) *Esp_Z* (1×10^6^ log-phase in 500 μl of LB media) and added the solution to trypanosome cultures maintained *in vitro*. This medium includes phenol red, which is a pH-sensitive dye that when in solution turns from red-pink to yellow as the quantity of acid in the environment increases. Addition of this HK *Esp_Z* extract immediately turned the Beck’s media yellow, and the pH measured at 5.8 (± 0.39). This value was significantly lower than trypanosome cultures that were supplemented with 500 μl of 1×10^6^ log-phase HK trypanosomes (pH 7.3 ± 0.28), HK *Sodalis* (pH 7.4 ± 0.39) or LB (*Esp_Z* growth media; pH 7.1 ± 0.28) or MM media (*Sodalis* growth media; pH 7.2± 0.29) alone (Figure 4A). We subsequently monitored trypanosome growth in cultures that received the above-mentioned supplements. We observed that trypanosomes failed to replicate in Beck’s medium that contained HK *Esp_Z* extracts, while trypanosomes multiplied in all of the other culture conditions (Figure 4B).

**Figure 4.**
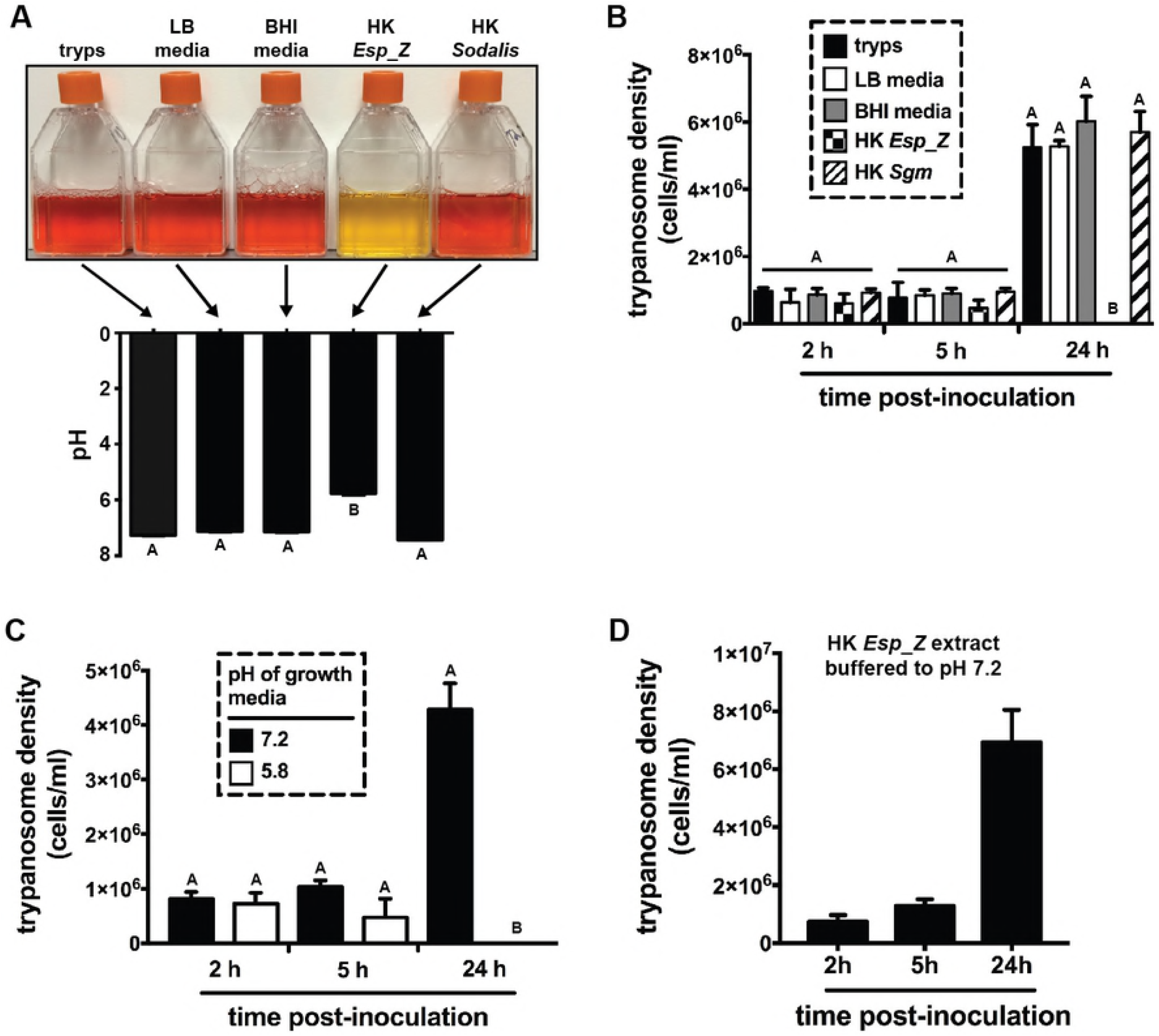
*Esp_Z* produces a low pH environment that is toxic to trypanosomes. (A) Early log phase trypanosomes (*T. b. brucei* YTAT 1.1), cultured in 10ml of Beck’s medium containing the pH sensitive dye phenol red, exposed to 1ml of heat treated LB media (*Esp_Z* culture medium), 1ml of heat treated BHI media (*Sodalis* culture medium), heat killed (HK) *Esp_Z* (5×10^6^ cells) in 1ml of LB media and HK *Sodalis* (5×10^6^ cells) in 1ml of BHI media. Controls are trypanosomes alone (tryps). All heated treatments and controls were allowed to cool to room temperature prior to adding them to the trypanosome cultures. Two hours post-treatment, culture pH was measured. HK *Esp_Z* significantly reduced the pH of the trypanosome culture (*p*<0.001). The experiment was repeated using 6 distinct clonal trypanosome populations (the image represents one of the six replicates). (B) Density of trypanosomes in culture 2h, 5h and 24h after addition of the treatments described in (A) above. At the 24h time point, all trypanosomes exposed to HK *Esp_Z* extracts were dead while those from the other groups were replicating similarly to controls. (C) Density of trypanosomes cultured in normal (pH 7.2) and artificially produced (via the addition of 0.1N HCl) acidic (pH 5.8) environments. Artificial acidic conditions kill all trypanosomes with 24 h. (D) The density of cultured trypanosomes exposed to HK *Esp_Z* extracts buffered to pH 7.2 (via the addition of 0.1N NaOH). The buffering treatment rescues parasite growth. In (A), (B) and (C), statistical significance was determined using a one-way ANOVA followed by Tukey’s HSD post-hoc analysis in (A), and a two-way ANOVA followed by Tukey’s HSD post-hoc analysis in (B) and (C). Different letters represents statistical significance between treatments and controls. In (B), (C) and (D), experiments were performed using 5 or 6 distinct clonal trypanosome populations.

Heat-killed *Esp_Z* extracts create an acidic environment when added to trypanosome cultures, and trypanosomes fail to replicate in this environment. These findings do not rule out the possibility that trypanosomes are capable of surviving *Esp_Z*-induced acidic conditions, and instead, some other unknown component of the medium [e.g., a bacterium-derived trypanocidal molecule(s)] exhibits toxic properties. To address this possibility, we monitored trypanosome growth in Beck’s medium, the pH of which was artificially decreased to 5.8 (the same as that achieved by adding HK *Esp_Z* extracts) via the addition of exogenous acid. Under these conditions trypanosomes failed to replicate (Figure 4C). Furthermore, when we buffered Beck’s medium containing HK *Esp_Z* extracts back up to pH 7.2, trypanosomes replicated normally (Figure 4D). Taken together, these data indicate that *Esp_Z* produces an acidic environment that is toxic to trypanosomes, thus impeding their growth *in vitro*.

### Esp_Z acidifies tsetse’s gut

We observed that trypanosomes are unable to multiply when cultured in medium supplemented with acidic *Esp_Z* extracts. Thus, we next investigated whether *Esp_Z* produces acid *in vivo* in tsetse’s gut. To do so we colonized teneral, aposymbiotic flies with either *Esp_Z* or *Sodalis*, and 5 days later fed them a meal containing 2.5% sucrose and 0.04% phenol red solubilized in water. Twenty-four hours later, midguts from a sample of flies (*n*=8 per group) were excised and plated on solid medium containing phenol red. *Gmm*^Apo/*Esp_Z*^ and *Gmm*^Apo/*Sgm*^ housed similar densities of the introduced bacteria (S1 Fig. B), and their respective mediums changed color to reflect corresponding pH shifts (S1 Fig. C). The remaining flies were dissected to expose their midgut *in situ*, and the color of the gut contents was visualized microscopically. We observed that the gut contents of *Gmm*^Apo/*Esp_Z*^ were yellow in color (Fig. 5), thus indicating that the environment had become acidified. Conversely, the gut contents of *Gmm*^Apo/*Sgm*^ individuals were red, which is similar to the more alkaline environment present in the gut of *Gmm*^WT^ tsetse (Fig. 5).

**Figure 5.**
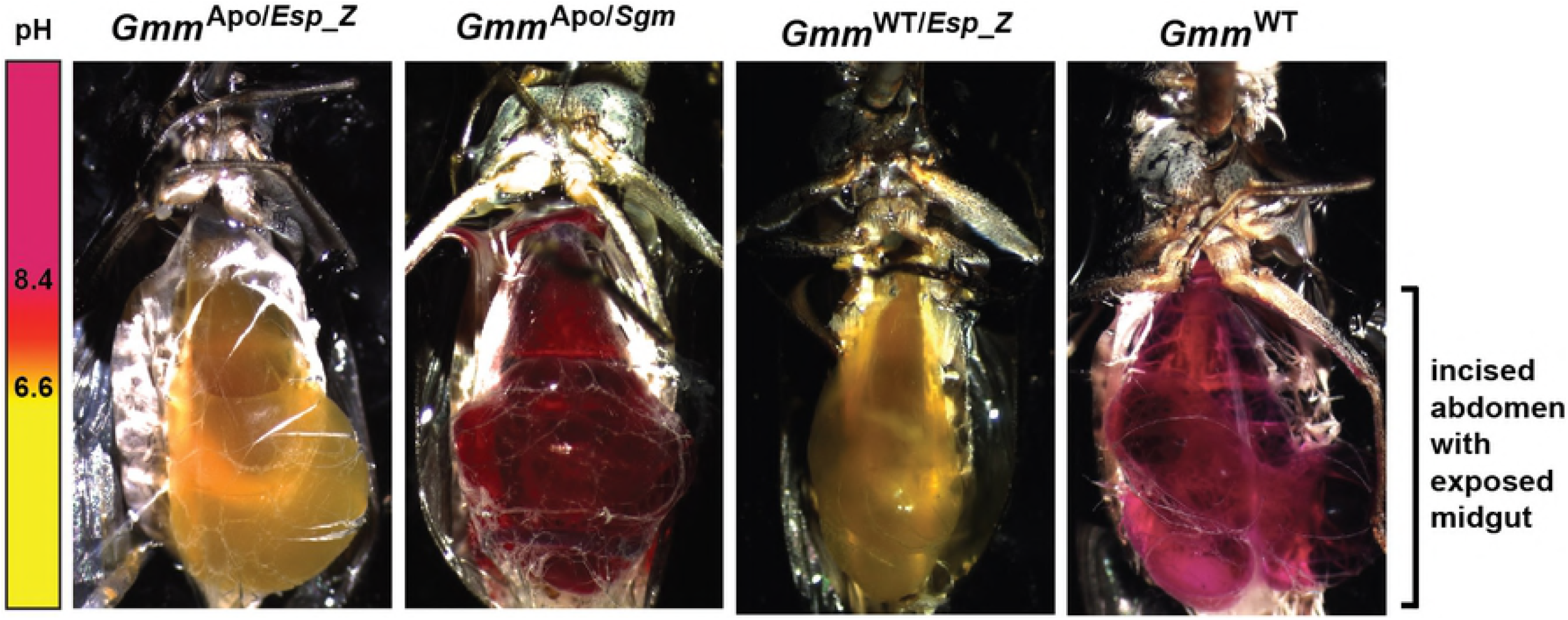
*Esp_Z* acidifies tsetse’s gut. Midgut pH of *Gmm*^Apo/*Esp_Z*^, *Gmm*^Apo/*Sgm*^, *Gmm*^WT/*Esp_Z*^ and *Gmm*^WT^ flies. Distinct groups of teneral *Gmm*^Apo^ flies were inoculated *per os* with either *Esp_Z* or *Sodalis*, while *Gmm*^WT^ individuals received *Esp_Z* (all flies received 5×10^4^ CFU of bacteria per ml of blood) or no bacteria. Five days post-inoculation, all individuals were offered a meal containing sucrose (2.5%) and phenol red (0.04%) dissolved in water. Twenty-four hours later, fly abdomens were excised and the color of the solution found within the midgut was observed. Each image represents one of five flies monitored for each treatment.

Finally, we investigated whether *Esp_Z* also produces acid in the gut *Gmm*^WT^ by inoculating teneral individuals with 1×10^3^ CFU of the bacterium (these flies were designated *Gmm*^WT/*Esp_Z*^). Five days later a cohort of *Gmm*^WT/*Esp_Z*^ females (these flies housed 1.27×10^6^ ± 8.6×10^4^ *Esp_Z* at this time point; S1 Fig. D), as well as age matched *Gmm*^WT^ controls, were fed a sugar meal containing phenol red (as described above) to observe gut pH. Similar to our results observed in *Gmm*^Apo/*Esp_Z*^, we observed that the gut of *Gmm*^WT/*Esp_Z*^ individuals was yellow, thus indicative of an acidified environment. Conversely, the gut environment of *Gmm*^WT^ was red and thus comparatively alkaline (Fig. 5). Thus, the presence of indigenous symbionts does not impede the ability of *Esp_Z* to acidify the gut of wild-type flies.

### Gmm^WT/Esp_Z^ are highly refractory to infection with trypanosomes and entomopathogenic bacteria

We hypothesized that exogenous microorganisms would be unable to successfully infect *Gmm*^WT/*Esp_Z*^ due to their acidified midgut environment. To test this hypothesis we first co-inoculated teneral *Gmm*^WT^ males with *Esp_Z* and *T. congolense* parasites. Two weeks post-challenge we observed no significant difference in the percentage of *Gmm*^WT/*Esp_Z*^ (15%) and control *Gmm*^WT^ (23%) that harbored trypanosome infections in their midguts (Fig. 6A). We next inoculated teneral *Gmm*^WT^ males with *Esp_Z* and then three days later (5 day old adults) challenged *Gmm*^WT/*Esp_Z*^ individuals with either *T. congolense* or *T. brucei* parasites (both of these parasite species are naturally transmitted by *G. m. morsitans;* Harley and Wilson, 1968; Moloo *et al*., 1992). Under these conditions we observed that *Gmm*^WT/*Esp_Z*^ males were significantly more refractory to infection with both parasite species (*T. congolense*, 6%; *T. brucei*, 32%) than were their age-matched *Gmm*^WT^ counterparts (*T. congolense*, 36%; *T. brucei*, 70%; Fig. 6B). Thus, tsetse must house an established *Esp_Z* infection in its gut at the time of trypanosome challenge in order to present a refractory phenotype. Finally, we found that trypanosome infected *Gmm*^WT/*Esp_Z*^ house similar densities of *Esp_Z* as do age-matched individuals that eliminated their trypanosome infection (S1 Fig. E), again indicating that exogenous *Esp_Z* appears to be resistant to tsetse’s trypanocidal immune response. Additionally, the presence of tsetse’s indigenous, enteric microbiota does not interfere with *Esp_Z* mediated obstruction of trypanosome infection establishment.

**Figure 6.**
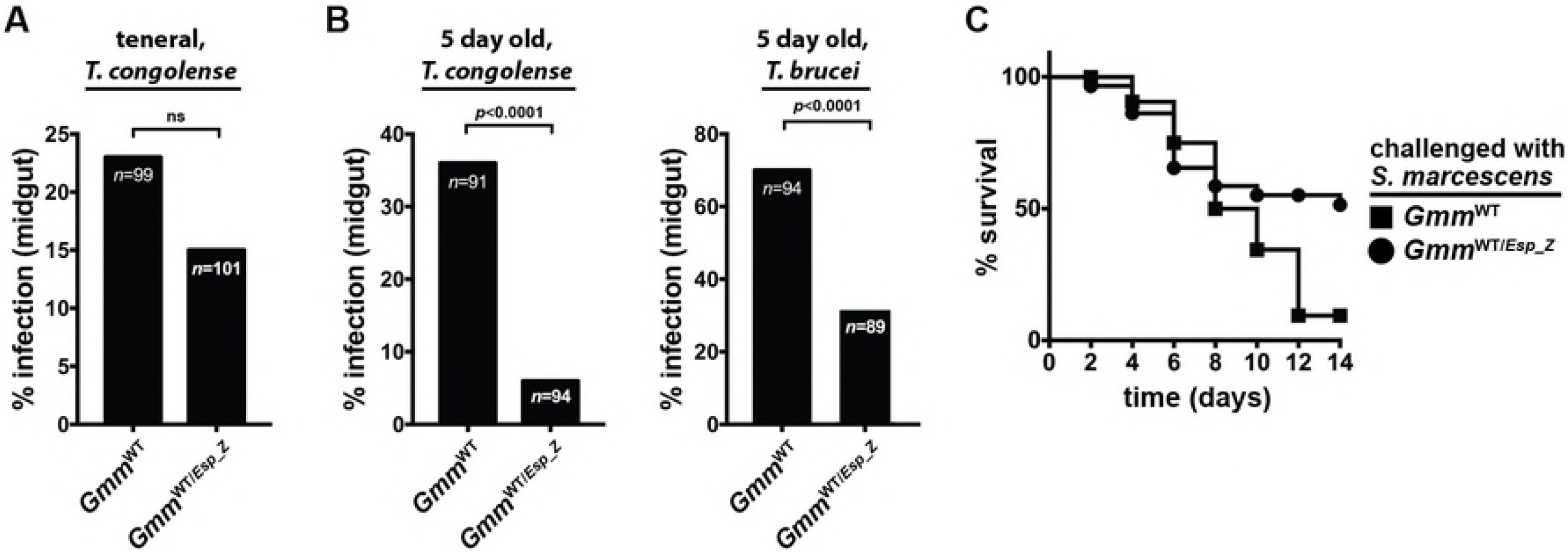
*Gmm*^WT/*Esp_Z*^ flies are significantly more refractory to infection with parasitic trypanosomes and entomopathogenic *S. marcescens*. (A) Percentage of *Gmm*^WT^ and *Gmm*^WT/*Esp_Z*^ flies infected with *T. congolense* 14 days after they were co-inoculated with *Esp_Z* and parasites in their first (teneral) blood meal. (B) Percentage of *Gmm*^WT^ and *Gmm*^WT/*Esp_Z*^ flies infected with *T. congolense* (left graph) and *T. brucei* (right graph). For these experiments *Gmm*^WT/*Esp_Z*^ flies housed the exogenous bacteria for fives prior to challenge with trypanosomes. In (A) and (B) Statistical analysis was performed using a GLM followed by multiple comparisons and Tukey contrasts, and different letters represents statistical significance between treatments and controls. (C) Kaplan-Meier plot depicting survival of *Gmm*^WT^ and *Gmm*^WT/*Esp_Z*^ flies following challenge with *S. marcescens*. Infection experiments were performed in triplicate, using 25 flies per replicate. *Gmm*^WT/*Esp_Z*^ flies were also significantly more refractory to *S. marcescens* infection than were their *Gmm*^WT^ counterparts (*p*=0.001; log-rank test).

Finally, we investigated whether *Esp_Z* also protects tsetse against infection with an entomopathogenic bacteria. To do so teneral *Gmm*^WT^ males were fed 1×10^3^ CFU of *Esp_Z*, and then three days later, the same dose of *Serratia marcescens* strain db11, which is highly virulent to wild-type tsetse (Weiss et al., 2014; Aksoy et al., 2016; Vigneron et al., 2018). Five day old *Gmm*^WT^ infected with the same dose of *S. marcescens* were used as controls. Fly survival following *Serratia* inoculation was monitored over a 14 day period in both fly groups. We observed that 51% and 9% of *Gmm*^WT/*Esp_Z*^ and *Gmm*^WT^ individuals, respectively, survived their infection with *S. marcescens* (Fig. 6C).

Taken together, our results detailed above indicate that wild-type tsetse present a parasite and entomopathogen refractory phenotype when they house an established *Esp_Z* infection in their gut. This phenotype like occurs because the acidified nature of the gut environment is inhospitable to the development of exogenous microbes.

### Esp_Z infection exerts a minimal fitness cost on tsetse

*Esp_Z* produces acid in tsetse’s gut such that the environment becomes inhospitable to trypanosomes. To address whether decreased midgut pH adversely impacts tsetse fitness, we quantified several fitness parameters in *Gmm*^WT/*Esp_Z*^ (a sample of these individuals housed 1.39×10^6^ ± 1.1×10^5^ *Esp_Z* at the time they were used for experimentation; S1 Fig. F).

We began by measuring midgut weight, which reflects over all digestive health. We observed no significant difference in midgut weight between 8 day old *Gmm*^WT/*Esp_Z*^ and *Gmm*^WT^ males (3.8 ± 1.1 mg and 3.2 ± 1.2 mg, respectively) and females (12.8 ± 1.8 mg and 13.0 ± 1.8 mg, respectively) 24 hrs post last blood meal acquisition (Fig. 7A). We next measured fecundity parameters in female and male *Gmm*^WT/*Esp_Z*^ and *Gmm*^WT^ to determine if stable infection with this bacterium would alter their reproductive capacity. We began by measuring gonotrophic cycle (GC) duration of *Gmm*^WT/*Esp_Z*^ and *Gmm*^WT^ females. The length of the 1^st^ GC was not significantly different between *Gmm*^WT/*Esp_Z*^ (24.0 ± 1.2 days) and *Gmm*^WT^ females (24.0 ± 0.9 days) (Fig. 7B). However, the 2^nd^ and 3^rd^ GCs of *Gmm*^WT/*Esp_Z*^ females (13.0 ± 1.1 and 14.0 ± 1.1 days, respectively) were significantly longer than those of their age-matched WT counterparts (14.0 ± 1.0 and 11.5 ± 1.1 days, respectively) (Fig. 7B). We also determined that pupal weight from all three GCs was similar between both fly groups (GC1, *Gmm*^WT/*Esp_Z*^ = 23.1 ± 1.5 mg, *Gmm*^WT^ = 22.9 ± 1.4 mg; GC2, *Gmm*^WT/*Esp_Z*^ = 23.4 ± 1.7 mg, *Gmm*^WT^ = 24.2 ± 1.8 mg; GC3, *Gmm*^WT/*Esp_Z*^ = 24.7 ± 1.7 mg, *Gmm*^WT^ = 24.4 ± 1.6 mg)(Fig. 7C). Thus, *Esp_Z* infection impacts the reproductive physiology of female tsetse by increasing GC duration and hence the number of offspring infected individuals are able to produce over the course of their lifespan. However, despite this effect, infection with this bacterium does not impact pupal weight.

**Figure 7.**
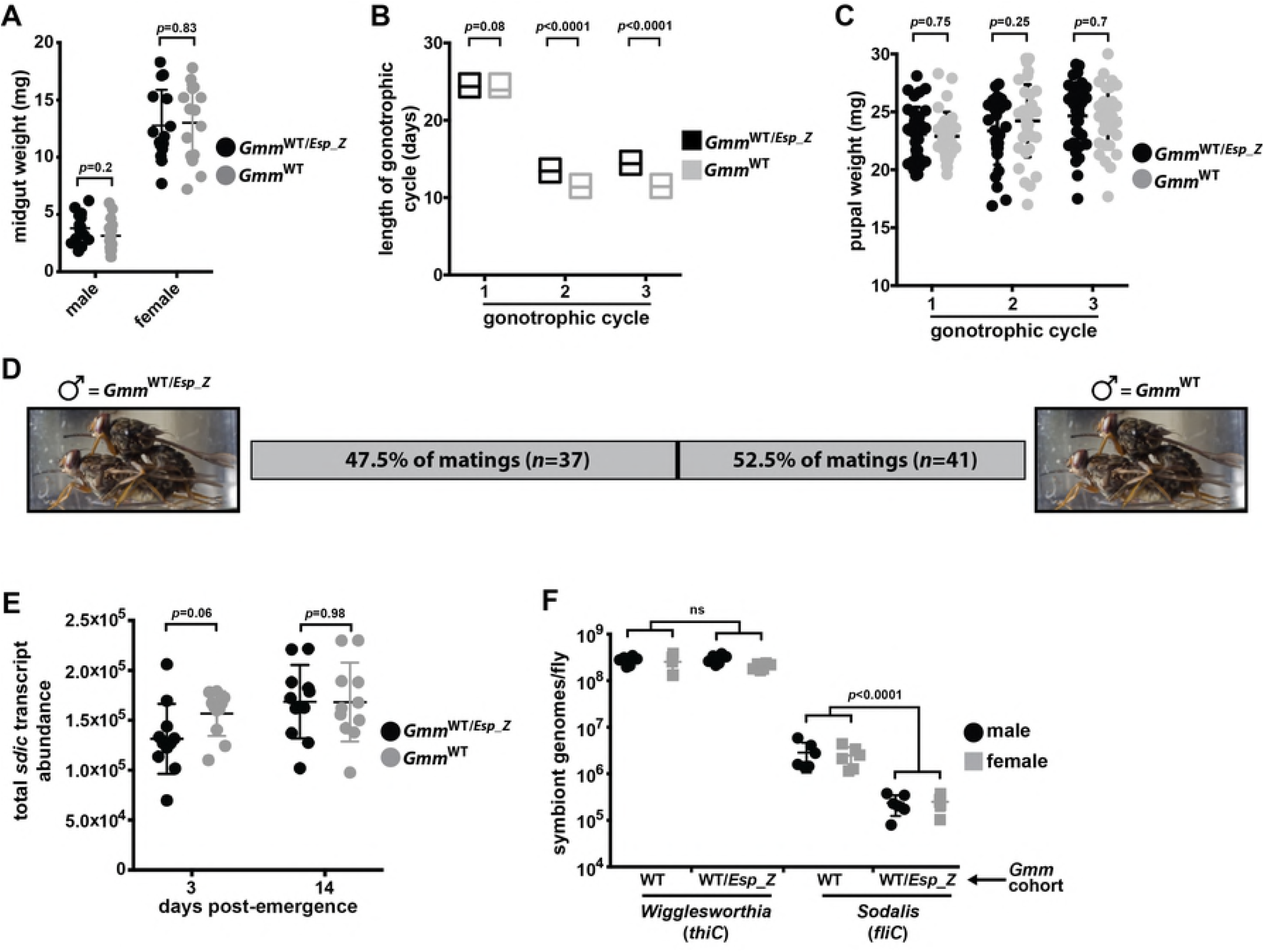
*Esp_Z* infection impacts specific tsetse fitness parameters. (A) Midgut weight, as an indicator of blood meal digestion proficiency, in six day old *Gmm*^WT/*Esp_Z*^ and *Gmm*^WT^ males and females (guts were weighed 24 h after the flies had consumed their last blood meal). Each point on the graph represents one individual, and statistical significance was determined via multiple t-tests. (B). Gonotrophic cycle (GC) length of *Gmm*^WT/*Esp_Z*^ and *Gmm*^WT^ females. Age-matched, pregnant females from each group (*n*=35 per group) were housed in individuals cages and monitored daily to observe frequency of pupal deposition. Statistical significance was determined via log-rank test. (C) Weight of pupae from *Gmm*^WT/*Esp_Z*^ and *Gmm*^WT^ females over three GCs. Each point on the graph represents one individual, and statistical significance was determined via multiple t-tests. (D) Mating competitiveness of *Gmm*^WT/*Esp_Z*^ compared to *Gmm*^WT^ males. Matings were setup in individual cages (*n*=80). Each cage housed a virgin female, to which one virgin *Gmm*^WT/*Esp_Z*^ and *Gmm*^WT^ male was added. Males and females were age-matched, and each had fed twice prior to exposure. Statistical significance was determined via Chi-squared test. (E) Sperm abundance in the reproductive tracts of three and 14 day old (fed twice) virgin *Gmm*^WT/*Esp_Z*^ and *Gmm*^WT^ males. Sperm quantity is a reflection of *sperm-specific dynein intermediate chain* (*sdic*) transcript abundance. Absolute *sdic* transcript abundance was determined by comparing experimental sample cycle threshold (C_t_) values to those derived from an *sdic* internal standard curve. Each point on the graph represents one individual, and statistical significance was determined via multiple t-tests. (F) Quantitation of *Wigglesworthia* and *Sodalis* in *Gmm*^WT/*Esp_Z*^ and *Gmm*^WT^ male and female tsetse. Abundance of symbiont specific gene transcripts (*Wigglesworthia, thiC; Sodalis, fliC*) was used as a proxy to quantify bacterial load. This was performed by comparing *thiC* and *fliC* cycle threshold (C_t_) values in *Gmm*^WT/*Esp_Z*^ and *Gmm*^WT^ flies to those derived from symbiont gene-specific internal standard curves. *Wigglesworthia* and *Sodalis* can be polyploid (Rio et al., 2006; Weiss et al., 2006), and as such, we normalized symbiont genome copy number to constitutively expressed tsetse *gapdh* copy number. Each point on the graph represents one individual, and statistical significance was determined via multiple t-tests.

We also investigated the effect of *Esp_Z* infection on the reproductive fitness of male tsetse by comparing the mating competitiveness of *Gmm*^WT/*Esp_Z*^ and *Gmm*^WT^ individuals. To do so we set up 80 individual cages, each of which contained one sexually mature virgin female. We subsequently placed one sexually mature *Gmm*^WT/*Esp_Z*^ and *Gmm*^WT^ male in each cage and monitored the arena to determine which of the two males successfully mated with the female. We observed that 47.5% of matings occurred between *Gmm*^WT/*Esp_Z*^ males and females (neither male mated with the female in two of the cages) (Fig. 7D), thus indicating that *Esp_Z* infection does not significantly alter male mating competitiveness. Next we compared the number of sperm present in three and 14 day old *Gmm*^WT/*Esp_Z*^ and *Gmm*^WT^ males by quantifying transcript abundance of *sperm-specific dynein intermediate chain* (*sdic*). The *Drosophila* homologue of this gene is transcribed exclusively in sperm cells (Nurminsky et al., 1998) and is used to quantify sperm abundance (Yeh et al., 2012). We observed no significant difference in *sdic* transcript abundance between three day old or 14 day old *Gmm*^WT/*Esp_Z*^ and *Gmm*^WT^ males (Fig. 7E).

Finally, we examined whether the presence of *Esp_Z* impacts the density of endosymbiotic *Wigglesworthia* and *Sodalis*. These measurements, which were taken at 14 days post-inoculation with *Esp_Z*, are important because tsetse’s microbiota impact many aspects of their host’s fitness, including fecundity and immune system development and function (Michalkova et al., 2014; Vigneron and Weiss, 2017). We observed that infection with *Esp_Z* did not significantly alter the density of tsetse’s midgut (bacteriome) population of obligate *Wigglesworthia* in *Gmm*^WT/*Esp_Z*^ males or females (Fig. 7F). Conversely, midguts from *Gmm*^WT/*Esp_Z*^ males and females housed significantly fewer *Sodalis* than did midguts from their *Gmm*^WT^ counterparts (Fig. 7F). Taken together, these data indicate that *Esp_Z* significantly impacts specific fitness parameters in both female and male flies.

## Discussion

Morbidity and mortality caused by vector-borne diseases currently inflicts a devastating socioeconomic burden on a significant percentage of the global population. To reduce this burden, novel disease control strategies that inhibit pathogen maturation within arthropod disease vectors require development. The enteric microbiota is being increasingly studied for use in this context, and one such novel strategy could employ the use of ‘probiotic’ bacteria, the presence of which would alter the physiology of the vector’s gut to make the environment inhospitable to pathogens. Herein we use the tsetse fly model system to highlight how an exogenous bacterium can be employed in this capacity to impede infection establishment of two pathogens in this insect disease vector. Specifically, we determined that *Esp_Z*, which is a bacterium found naturally in the gut of some *An. gambiae* populations, and directly kills *Plasmodium* by producing anti-parasitic ROIs (Cirimotich et al., 2011), can stably colonize tsetse’s gut for at least 28 days. When the bacterium is present in this niche, tsetse are significantly more refractory to infection with parasitic African trypanosomes and entomopathogenic *S. marcescens* than are flies that house only their indigenous microbiota. *Esp_Z* creates this inimical environment by acidifying tsetse’s gut such that trypanosomes and *S. marcescens*, which are sensitive to these conditions, are no longer able to successfully infect the fly. While infection with *Esp_Z* exerts only a negligible effect on tsetse’s reproductive fitness, the bacterium’s presence does reduce the density of endosymbiotic *Sodalis*. Cumulatively, our findings suggest that *Esp_Z* could be used in a natural setting to artificially reduce disease transmission by this arthropod vector.

Herein we demonstrate that *Esp_Z* is able to stably colonize tsetse’s gut, which is an outcome that likely results at least in part from the bacterium’s resistance to antimicrobial PGRP-LB. This protein is constitutively produced in the fly’s midgut and directly kills trypanosomes (Wang et al., 2012) and *E. coli* K12 (Fig. 2 in this study, and Wang et al., 2012). *Esp_Z* resistance to tsetse PGRP-LB may reflect one of many bacterial adaptations that result from residing within the immunologically hostile environment of the insect midgut. While the specific physiological mechanism(s) that *Esp_Z* uses to facilitate its colonization of tsetse’s midgut are currently unknown, the bacterium survives for prolonged periods within the gut of *An. gambiae* in part by increasing expression of genes that encode a type III secretion system apparatus protein as well as glutathione S-transferase and oxidoreductase (Dennision et al., 2016). Type III secretion system proteins can facilitate bacterial penetrance into host cells (Dale and Moran, 2006) and be involved in subversion of host immunity (Raymond et al., 2013), while the latter two proteins are antioxidant pathway components that mediate redox homeostasis in oxidatively stressful environments such as the insect midgut (Ketterman et al., 2011; Pedrini et al., 2015). *Esp_Z* may employ similar mechanisms to survive in tsetse’s immunologically hostile gut. *Sodalis* is also resistant to tsetse antimicrobial immune response (Hu and Aksoy, 2005; Wang et al., 2013), which may be the result of structural adaptations present in exposed bacterial surface coat molecules, including lipopolysaccharide (Toh et al., 2006) and outer membrane protein A (Weiss et al., 2008). Furthermore, *Sodalis* enters into host cells through the use of a type III secretion system (Dale et al., 2001), which may further protect the bacterium from tsetse’s immunologically hostile midgut environment. Likewise, similar mechanisms may facilitate *Esp_Z* survival in this niche. Additionally, *Esp_Z*, like *Sodalis* (Maltz et al., 2012), may reside extracellularly in the endoperitrophic space of tsetse’s midgut. In this position, the fly’s peritrophic matrix barrier would physically separate the bacteria from immunocompetent epithelial cells, thus reducing their exposure to harmful antimicrobial responses.

Microbes can alter their environment such that it either favors or hinders its own prosperity as well as the prosperity of other resident organisms (Ratzke et al., 2018a). Depending on specific physiological circumstances, these effects can reflect the consumption of resources and/or the production of beneficial or harmful metabolic byproducts (Celiker and Gore, 2013; Ratzke et al., 2018b). Tsetse’s sole energy source, vertebrate blood, is rich in glucose. Many bacterial taxa, including *Enterobacter* spp., ferment this sugar, thus producing hydrogen that acidifies their environment (Ratzke et al., 2018b, Goldford et al., 2018). The acidic environment present in tsetse’s gut when *Esp_Z* resides stably in the tissue likely results at least in part from the bacterium’s utilization of blood glucose as an energy source. The pH in the gut of insect vectors, including tsetse flies (Liniger et al., 2003), sand flies (Rosenzweig et al., 2007) and mosquitoes (del Pilar Corena et al., 2005) is normally alkaline, and the parasites they transmit, as well as other enteric microbes, are adapted to survive in this environment. Correspondingly, our results indicate that *Esp_Z* induced conditions in tsetse’s gut detrimentally impact not only trypanosomes but also other enteric microbes including entomopathogenic *S. marcescens* and symbiotic *Sodalis. Esp_Z* mediated suppression of *S. marcescens*, or any other pathogen, would have obvious benefits to the fly. However, dysbiosis of tsetse’s facultative and commensal enteric microbiota could impact the fly’s overall fitness and/or vector competency. For example, reducing *Sodalis* density significantly decreases tsetse longevity (Dale and Welburn, 2001). This may prove beneficial because flies with a reduced life span could perish before trypanosomes are able to complete their 20-30 extrinsic incubation period (Aksoy et al., 2001). A reduction in *Sodalis* density could be further beneficial because tsetse that house relatively low densities of the bacterium are less likely to be infected with trypanosomes than are individuals that house more of the symbiont (Welburn et al., 1993; Dale and Welburn, 2001; Farikou et al., 2010; Soumana et al., 2013; Griffith et al., 2018). Thus, the trypanosome refractory phenotype presented by *Esp_Z* colonized tsetse may result in part from, or be enhanced by, the fact that they contain fewer *Sodalis*. Finally, the midgut of wild tsetse is colonized by a transient population of environmentally acquired bacteria (Wang et al., 2013). The contribution of these bacteria to tsetse’s physiology has not been characterized, and as such, interference with this microbial population could further alter the fly’s physiological homeostasis. To the contrary, the environmentally acquired microbiota could out-compete *Esp_Z* that reside in tsetse’s gut, or could prevent the bacterium from acidifying the environment. Future studies are required to elucidate microbe-microbe interactions in the gut *Esp_Z* colonized flies after their release into the field.

Reducing the incidence of African trypanosomiases has to date been achieved largely by controlling the size of tsetse populations. This process is currently accomplished by employing area wide integrated pest management (AW-IPM) strategies that make use of insecticides, traps and sterile insect technique (SIT) (Vreyson et al., 2013; Percoma et al., 2018). SIT involves sequentially releasing a large number of sterilized males (achieved by irradiating pupae) into the target environment. These males reproductively outcompete wild males for female mates, and the population size drops significantly, or the fly is completely eradicated (McGraw and O’Neill, 2013). The efficacy of SIT as a means of controlling tsetse populations is well exemplified on Unguja Island (the large island of the Zanzibar archipelago), where the technique was used to eradicate *G. austeni*, the main vector of trypanosomes that cause animal African trypanosomiasis in that locale (Vreysen et al., 2000). One shortcoming of this procedure is that releasing large numbers of sterile males significantly increases the population of potential disease vectors in that environment (male tsetse also feed exclusively on vertebrate blood). One way to overcome this obstacle is to release sterilized males that present enhanced refractoriness to parasite infection. This outcome is currently achieved by feeding the sterilized males twice with the drug isometamidium chloride prior to their release (Bouyer, 2008). However, treated flies are not 100% resistant to infection (Bouyer, 2008), and the risk exists that the parasite will eventually develop resistance to the drug. Our data presented herein indicate that inoculating sterilized males with *Esp_Z* prior to their release would serve as an alternative, or supplemental, means of making the flies resistant to infection. Specifically, *Gmm*^WT/*Esp_Z*^ males (and females) are significantly more refractory to infection with trypanosomes than are their wild-type counterparts. This finding implies that sterilized, *Gmm*^WT/*Esp_Z*^ individuals would be relatively poor vectors of disease-causing trypanosomes and thus safer to release than sterilized males that do not house this bacterium. Furthermore, our preliminary analyses suggest that *Esp_Z* infection does not compromise the mating competitiveness nor sperm abundance of male tsetse, thus suggesting that *Gmm*^WT/*Esp_Z*^ individuals would be as successful as their wild counterparts at locating females and engaging in viable matings. Finally, male tsetse could be irradiated as pupae or teneral adults (de Beer et al., 2017), prior to colonization with *Esp_Z*, thus eliminating the possibility that this treatment could detrimentally impact the bacterium’s fitness and thus its effect on fly vector competency. These characteristics provide preliminary evidence that releasing sterilized, *Esp_Z* infected male tsetse as part of an AW-IPM program would significantly reduce the capacity of these flies to transmit disease.

In conclusion, data presented in this study indicates that *Esp_Z* could effectively complement currently used AW-IPM programs aimed at reducing or eliminating tsetse populations by inhibiting trypanosome infection establishment in the fly’s gut. However, the complex relationship between tsetse, its indigenous (endosymbionts) and exogenous (trypanosomes and environmentally acquired microorganisms) microbiota, and *Esp_Z* must be studied in more detail before the bacterium is used in this capacity. Of particular importance are studies aimed at determining whether *Esp_Z* presents trypanocidal activity in other epidemiologically important tsetse species (e.g., *G. fuscipes*). Furthermore, field-based studies would shed light on how the ecology of tsetse’s natural environment influences the overall efficacy of the system.

## Materials and Methods

### Ethical Consideration

This work was carried out in strict accordance with the recommendations in the Office of Laboratory Animal Welfare at the National Institutes of Health and the Yale University Institutional Animal Care and Use Committee. The experimental protocol was reviewed and approved by the Yale University Institutional Animal Care and Use Committee (Protocol 2011-07266).

### Tsetse, bacteria and trypanosomes

Tsetse flies (*Glossina morsitans morsitans*) were maintained in Yale University’s insectary at 24°C with 55% relative humidity. Flies received defibrinated bovine blood through an artificial membrane feeding system every 48 h. Aposymbiotic tsetse (*Gmm*^Apo^) were generated and maintained as described previously (Weiss et al., 2012). Throughout the manuscript, flies referred to as ‘teneral’ were unfed adults recently eclosed (≤ 24h) from their pupal case. All tsetse lines used in this study are described in S1 Table.

*Sodalis* were isolated from tsetse pupae as described previously (Dale and Maudlin, 1999), and subsequently maintained in liquid brain heart infusion (BHI) media. When necessary, *Sodalis* were plated on either Brain Heart Infusion agar supplemented with 10% defibrinated bovine blood (BHIB) or Mitsuhashi-Maramorosch (MM)-agar plates. *Enterobacter* sp Z (*Esp_Z;* isolated from the gut of the mosquito, *Anopheles gambiae*) (Cirimotich et al., 2011) and *Serratia marcescens* (strain db11; isolated from a moribund *Drosophila* sp.) (Nehme et al., 2007) were grown in liquid LB media or on LB-agar plates at 30°C.

*In vivo Trypanosoma congolense* and *T. brucei brucei* (YTAT 1.1) were expanded in rats, and harvested from infected blood at peak parasitemia. Rat blood containing blood stream form (BSF) parasites was aliquoted and cryopreserved for subsequent tsetse challenge experiments.

### Recombinant PGRP-LB antibacterial assays

Antibacterial activity of recombinant (rec) PGRP-LB was determined as described previously (Wang et al., 2012), with minor modification. Specifically, recPGRP-LB was added (10 μg/ml of media) to early log-phase (OD = 0.2-0.4) cultures of *Esp_Z, Sodalis* and *E. coli*. Controls consisted of bacterial cultures exposed to bovine serum albumin. Using a plate-based quantification assay (Maltz et al., 2012), *E. coli* and *Esp_Z* density was subsequently measured 1 hr. later, while *Sodalis* density was measured 24 hr. later. Results are presented as % of initial inoculum, which was determined by dividing the number of bacterial CFU present after treatment with recPGRP-LB by the number of CFU present prior to inoculation.

### Microbial infection assays

*Per os* bacterial challenge of wild-type (*Gmm*^WT^) and *Gmm*^Apo^ flies was performed by feeding teneral adults a heat inactivated (HI; 56°C for 1 hr) blood meal inoculated with 5×10^4^ colony forming units (CFU) of each bacterial strain per ml of blood. Because tsetse flies consume approximately 20 μl of blood per feeding, each fly is inoculated with 1×10^3^ bacterial cells. *Gmm*^Apo^ flies colonized with either *Sodalis* or *Esp_Z* are designated *Gmm*^Apo/*Sgm*^ and *Gmm*^Apo/*Esp_Z*^, respectively, and *Gmm*^WT^ flies colonized with *Esp_Z* are designated *Gmm*^WT/*Esp_Z*^. For all experiments that employed tsetse flies inoculated with either *Sodalis* or *Esp_Z*, bacterial midgut density was determined by homogenizing microscopically dissected gut tissue in 0.85% NaCl and serially diluting and plating the samples on LB-agar (*E. coli, Esp_Z* and *Serratia*) or BHIB or MM (*Sodalis*) plates supplemented with antibiotics. CFU per plate were counted manually, and counts are presented in the corresponding Results subsections.

For trypanosome infections, all flies received infectious blood meals containing 1×10^6^/mL BSF *T. congolense* or *T. b. brucei* parasites. *Gmm*^Apo^, *Gmm*^Apo/*Sgm*^ and *Gmm*^Apo/*Esp_Z*^ were challenged as eight day old adults (3^rd^ blood meal), while *Gmm*^WT^ and *Gmm*^WT/*Esp_Z*^ flies were challenged as five day old adults (2^nd^ blood meals). For *Esp_Z*/trypanosome co-infection experiments, distinct groups of mature *Gmm*^WT^ individuals were inoculated with 1×10^6^/mL BSF *T. congolense* parasites and 5×10^4^ CFU/ml of *Esp_Z*. Two weeks post-trypanosome challenge, all flies were dissected and their midguts microscopically examined to determine parasite infection status.

### Detection and inhibition of tsetse reactive oxygen intermediates

*Esp_Z* and *Sodalis* cultures were grown to mid-log phase (OD = 0.25), and cell-free supernatants were generated via centrifugation. Hydrogen peroxide (H_2_O_2_) concentrations in bacterial culture supernatants were determined using an Amplex Red Hydrogen Peroxide/Peroxidase assay kit according to the manufacturer’s (Invitrogen) protocol. In brief, supernatants were incubated for 30 min. with the assay reagent, and resulting fluorescence units were quantified using a Bio-Tek plate reader.

Antioxidants were used to inhibit tsetse ROI activity *in vivo*. The assay used was similar to those described previously (MacLeod et al., 2007; Cirimotich et al., 2011; Vigneron et al., 2018). In brief, treated flies were offered a blood meal inoculated with trypanosomes [1×10^6^/mL BSF *T. b. brucei* (YTAT 1.1)] and supplemented with vitamin C (10mM) or cysteine (10μM). All subsequent meals also contained antioxidant supplements.

### Determination of bacterial acid production in vitro

*Sodalis* and *Esp_Z* were grown in their respective liquid media to an O.D. of 1.0. Subsequently, 5×10^6^ cells (this value represents the approximate maximum density to which these bacteria grow in tsetse’s gut; see Fig. 1A) were diluted to a volume of 1 ml (again in respective liquid media) and heat-killed (80°C for 1.5 hr). Conditioned media containing dead cells was added to early log growth phase *T. b. brucei* YTAT 1.1 grown in a Beck’s medium (GE Hyclone), which contains phenol red. When in solution this compound serves as a pH-sensitive colorimetric indicator that changes from pink-red to yellow as environmental pH drops. Other treatment groups were inoculated with 1 ml of heated, clean LB (*Esp_Z* growth medium) or clean BHI (*Sodalis* growth medium), while the control group consisted of trypanosomes alone. Two hours after exposing *T. b. brucei* to treatment conditions, cultures were assayed to determine pH using a Mettler Toledo pH meter. The pH of trypanosome containing Beck’s medium was experimentally reduced (to pH 5.8) via the addition of 0.1N HCl, while HK *Esp_Z* extracts were buffered to pH 7.2 via the addition of 0.1N NaOH. Trypanosome density in all treatment and control groups was determined at 2, 5 and 24 hour time points by counting live parasites using a Brite-Line hemocytometer.

### Determination of bacterial acid production in vivo

Microbial regulation of pH in tsetse’s midgut was determined by feeding teneral *Gmm*^Apo^ flies a HI blood meal inoculated with either *Sodalis* or *Esp_Z* (5×10^4^ CFU/ml of blood). Additionally, teneral *Gmm*^WT^ flies received the same quantity of *Esp_Z*. Five days post-bacterial challenge, colonized individuals were administered a meal composed of sucrose (2.5%) and phenol red (0.04%) solubilized in water. Twenty-four hours later, the color of the solution contained in the midgut was determined by incising the fly abdomen and observing the intact gut using a dissecting microscope (Zeiss Discovery) equipped with a digital camera (Zeiss AxioCam MRc 5). Remaining flies were dissected and their guts were harvested, homogenized in 0.85% NaCl, serially diluted and plated onto MM-agar plates (prepared as described in Weiss et al., 2008) supplemented with phenol red (0.025 g/L) and sucrose (a 2.5% sucrose solution was spread onto plates immediately prior to applying tsetse gut extracts). CFU per plate was counted manually, and the growth medium was monitored to observe pH-induced changes in color.

### Fitness assays

For all fitness assays, *Gmm*^WT^ teneral females and males were infected with *Esp_Z* during their first blood meal. To determine midgut weight, midguts were dissected from 9 day old *Gmm*^WT/*Esp_Z*^ and *Gmm*^WT^ females and males (24 h after consuming their last blood meal) and weighed using a Mettler Toledo (AL104) balance. The effect of *Esp_Z* infection on female fecundity was measured by quantifying the length of three gonotrophic cycles (GC) and by weighing pupal offspring. To measure GC length, *Gmm*^WT/*Esp_Z*^ and *Gmm*^WT^ females were mated as 5 day old adults and thereafter maintained in individual cages. All females were monitored daily to determine when they deposited larvae, and all deposited larvae were weighed.

The effect of *Esp_Z* infection on male reproductive fitness was measured by quantifying the mating competitiveness and sperm abundance of individuals that housed the bacterium versus those that did not. Mating competitiveness assays were performed in individual cages, each of which housed one 5 day old virgin female (fed twice). Subsequently, one age-matched *Gmm*^WT/*Esp_Z*^ and *Gmm*^WT^ male (also fed twice) was added to each cage. These males were distinguished from one another by removing the proximal tarsus of the right foreleg from one of the individuals. The arena was observed until one of the males had successfully mounted the female, at which point the cage was submerged in ice and the free male identified. To eliminate any bias associated with removal of the foreleg tarsus, the experiment was repeated twice (*n*=40 cages per experiment), each time with either *Gmm*^WT/*Esp_Z*^ or *Gmm*^WT^ males receiving the distinguishing procedure. Sperm abundance was measured by RT-qPCR quantification of *sperm-specific dynein intermediate chain* (*sdic*) expression in the reproductive tracts of three and 14 day old (fed twice) virgin *Gmm*^WT/*Esp_Z*^ and *Gmm*^WT^ males. Absolute *sdic* transcript abundance was determined by comparing experimental sample cycle threshold (C_t_) values to those derived from an *sdic* internal standard curve.

*Sodalis fliC* and *Wigglesworthia thiC* gene specific primers were used to quantify the absolute abundance of these bacteria. This was performed by comparing *Sodalis fliC* and *Wigglesworthia thiC* cycle threshold (C_t_) values in *Gmm*^WT/*Esp_Z*^ and *Gmm*^WT^ females and males to those derived from symbiont gene-specific internal standard curves. Because *Wigglesworthia* and *Sodalis* can be polyploid (Rio et al., 2006; Weiss et al., 2006), we normalized symbiont genome copy number to constitutively expressed tsetse *gapdh* copy number. All RT-qPCR primers are listed in S2 Table. All RT-qPCR assays were carried out in duplicate, and replicates were averaged for each sample. Negative controls were included in all amplification reactions.

### Statistical analyses

For trypanosome infection experiments, statistical analyses were carried out using the R software for macOS (version 3.3.2) or GraphPad Prism(v.6). A generalized linear model (GLM) was generated using binomial distribution with a logit transformation of the data. The binary infection status (infected or recovered) was analyzed as a function of the bacterium used to colonized the insects (or its absence). For experiments requiring a pairwise comparison, we performed a Wald test on the individual regression parameter (nature of the bacterium used to colonize) to test its statistical difference. For experiments requiring multiple comparisons, multiple pairwise tests were generated using Tukey contrasts on the generalized linear model (GLM) using glht() function of “multcomp” package in R. Details of the statistical tests described above are indicated in S1 Dataset. All statistical tests used, and statistical significance between treatments, and treatments and controls, are indicated on the figures or in their corresponding legends. All samples sizes are provided in corresponding figure legends or are indicated graphically as points on dot plots.

## Acknowledgements

We thank Dr. George Dimopoulos (Department of Molecular Microbiology and Immunology, Johns Hopkins Bloomberg School of Public Health) for generously sharing *Esp_Z*. We thank members of the Aksoy lab for providing critical review of the manuscript. We thank the International Atomic Energy Association (IAEA), under the auspices of a Coordinated Research Project entitled ‘Enhancing tsetse fly refractoriness to trypanosome infection’, and Dr. Peter Takac (Institute of Zoology Slovak Academy of Science), for providing *G. morsitans* pupae used in this study.

## Supporting information

**S1 Fig. Density [colony forming units (CFU) per fly gut] of exogenous *Esp_Z* and *Sodalis* in the gut of experimental flies, and *Esp_Z* acid production *in vitro*.** *Esp_Z* and *Sodalis* CFU/gut in *Gmm*^Apo/*Sgm*^ and *Gmm*^Apo/*Esp_Z*^ flies prior to (A) challenge with trypanosomes and (B) measuring gut pH *in vivo*. (C) Guts form *Gmm*^Apo/*Sgm*^ and *Gmm*^Apo/*Esp_Z*^ flies homogenized and plates onto MM-agar plates supplemented with phenol red (0.025 g/L) and sucrose (a 2.5% sucrose solution was spread onto plates immediately prior to applying gut extracts). Plate color reflects bacteria induced changes in pH relative to the empty control. (D) *Esp_Z* density in the gut of *Gmm*^WT/*Esp_Z*^ flies prior to measuring gut pH *in vivo*. (E) *Esp_Z* density in the gut of trypanosome infected (TI) and trypanosome refractory (TR) *Gmm*^WT/*Esp_Z*^ flies. Measurements were taken at the time infection status was determined (14 days post-challenge). (F) *Esp_Z* density in the gut of a random sample of *Gmm*^WT/*Esp_Z*^ flies used to determine the bacterium’s impact of tsetse fitness parameters.

## References

Dale C, Moran NA. Molecular interactions between bacterial symbionts and their hosts. Cell. 2006;126: 453–465.

Goldford JE, Lu N, Bajić D, Estrela S, Tikhonov M, Sanchez-Gorostiaga A, et al. Emergent simplicity in microbial community assembly. Science 2018;361: 469–474.

Bouyer J. Does isometamidium chloride treatment protect tsetse flies from trypanosome infections during SIT campaigns? Med Vet Entomol. 2008;22: 140–143.

Aksoy S, O’Neill SL, Maudlin I, Dale C, Robinson AS. Prospects for control of African trypanosomiasis by tsetse vector manipulation. Trends Parasitol. 2001;17: 29–35.

Moloo SK, Sabwa CL, Kabata JM. Vector competence of *Glossina pallidipes* and *G. morsitans centralis* for *Trypanosoma vivax, T. congolense and T. b. brucei*. Acta Trop. 1992;51: 271–280.

Harley JM, Wilson AJ. Comparison between *Glossina morsitans, G. pallidipes* and *G. fuscipes* as vectors of trypanosomes of the *Trypanosoma congolense* group: the proportions infected experimentally and the numbers of infective organisms extruded during feeding. Ann Trop Med Parasitol. 1968;62: 178–187.

Dey R, Joshi AB, Oliveira F, Pereira L, Guimarães-Costa AB, Serafim TD, et al. Gut microbes egested during bites of infected sand flies augment severity of Leishmaniasis via inflammasome-derived IL-1β. Cell Host Microbe 2018;23: 134–143.

Soumana IH, Simo G, Njiokou F, Tchicaya B, Abd-Alla AM, Cuny G, et al. The bacterial flora of tsetse fly midgut and its effect on trypanosome transmission. J Invertebr Pathol. 2013;112 Suppl: S89–93.

Farikou O, Njiokou F, Mbida Mbida JA, Njitchouang GR, Djeunga HN, Asonganyi T, Et al. Tripartite interactions between tsetse flies, *Sodalis glossinidius* and trypanosomes--an epidemiological approach in two historical human African trypanosomiasis foci in Cameroon. Infect Genet Evol. 2010;10: 115–121.

Balmand S, Lohs C, Aksoy S, Heddi A. Tissue distribution and transmission routes for the tsetse fly endosymbionts. J Invertebr Pathol. 2013;112 Suppl: S116–122.

Pedrini N, Ortiz-Urquiza A, Huarte-Bonnet C, Fan Y, Juárez MP, Keyhani NO. Tenebrionid secretions and a fungal benzoquinone oxidoreductase form competing components of an arms race between a host and pathogen. Proc Natl Acad Sci USA. 2015;112: E3651–3660.

Ketterman AJ, Saisawang C, Wongsantichon J. Insect glutathione transferases. Drug Metab Rev. 2011;43: 253–265

Raymond B, Young JC, Pallett M, Endres RG, Clements A, Frankel G. Subversion of trafficking, apoptosis, and innate immunity by type III secretion system effectors. Trends Microbiol. 2013;21: 430–441.

Dale C, Young SA, Haydon DT, Welburn SC. The insect endosymbiont Sodalis glossinidius utilizes a type III secretion system for cell invasion. Proc Natl Acad Sci USA. 2001;98: 1883–1888.

Toh H, Weiss BL, Perkin SA, Yamashita A, Oshima K, Hattori M, et al. Massive genome erosion and functional adaptations provide insights into the symbiotic lifestyle of *Sodalis glossinidius* in the tsetse host. Genome Res. 2006;16: 149–156.

Hu Y, Aksoy S. An antimicrobial peptide with trypanocidal activity characterized from *Glossina morsitans morsitans*. Insect Biochem Mol Biol. 2005;35: 105–115.

Weiss BL, Wu Y, Schwank JJ, Tolwinski NS, Aksoy S. An insect symbiosis is influenced by bacterium-specific polymorphisms in outer-membrane protein A. Proc Natl Acad Sci USA. 2008;105): 15088–15093.

MacLeod ET, Maudlin I, Darby AC, Welburn SC. Antioxidants promote establishment of trypanosome infections in tsetse. Parasitology 2007;134: 827–831.

Abd-Alla AM, Bergoin M, Parker AG, Maniania NK, Vlak JM, Bourtzis K, Boucias DG, et al. Improving Sterile Insect Technique (SIT) for tsetse flies through research on their symbionts and pathogens. J Invertebr Pathol. 2013;112 Suppl: S2–10.

Van Den Abbeele J, Bourtzis K, Weiss B, Cordón-Rosales C, Miller W, Abd-Alla AM, et al. Enhancing tsetse fly refractoriness to trypanosome infection--a new IAEA coordinated research project. J Invertebr Pathol. 2013;112 Suppl: S142–147.

Aksoy S, Weiss B, Attardo G. Paratransgenesis applied for control of tsetse transmitted sleeping sickness. Adv Exp Med Biol. 2008;627: 35–48.

De Vooght L, Caljon G, De Ridder K, Van Den Abbeele J. Delivery of a functional anti-trypanosome Nanobody in different tsetse fly tissues via a bacterial symbiont, *Sodalis glossinidius*. Microb Cell Fact. 2014;13: 156.

De Vooght L, Caljon G, Van Hees J, Van Den Abbeele J. Paternal transmission of a secondary symbiont during mating in the viviparous tsetse fly. Mol Biol Evol. 2015;32: 1977–1980.

Dale C, Welburn SC. The endosymbionts of tsetse flies: manipulating host-parasite interactions. Int J Parasitol. 2001;31: 628–631.

Celiker H, Gore J. Cellular cooperation: insights from microbes. Trends Cell Biol. 2013;23: 9–15.

del Pilar Corena M, VanEkeris L, Salazar MI, Bowers D, Fiedler MM, Silverman D, et al. Carbonic anhydrase in the adult mosquito midgut. J Exp Biol. 2005;208: 3263–3273.

Rosenzweig D, Smith D, Opperdoes F, Stern S, Olafson RW, Zilberstein D. Retooling Leishmania metabolism: from sand fly gut to human macrophage. FASEB J. 2008;22: 590–602.

Liniger M, Acosta-Serrano A, Van Den Abbeele J, Kunz Renggli C, Brun R, Englund PT, et al. Cleavage of trypanosome surface glycoproteins by alkaline trypsin-like enzyme(s) in the midgut of *Glossina morsitans*. Int J Parasitol. 2003;33: 1319–1328.

Percoma L, Sow A, Pagabeleguem S, Dicko AH, Serdebéogo O, Ouédraogo M, et al. Impact of an integrated control campaign on tsetse populations in Burkina Faso. Parasit Vectors. 2018;11: 270.

Ratzke C, Denk J, Gore J. Ecological suicide in microbes. Nat Ecol Evol. 2018a;2: 867–872.

Ratzke C, Gore J. Modifying and reacting to the environmental pH can drive bacterial interactions. PLoS Biol. 2018b;16: e2004248.

Yeh SD, Do T, Chan C, Cordova A, Carranza F, Yamamoto EA, et al. Functional evidence that a recently evolved *Drosophila* sperm-specific gene boosts sperm competition. Proc Natl Acad Sci USA. 2012;109: 2043–2048.

Coon KL, Vogel KJ, Brown MR, Strand MR. Mosquitoes rely on their gut microbiota for development. Mol Ecol. 2014;23: 2727–2739.

Coon KL, Brown MR, Strand MR. Mosquitoes host communities of bacteria that are essential for development but vary greatly between local habitats. Mol Ecol. 2016;25: 5806–5826.

Bahia AC, Dong Y, Blumberg BJ, Mlambo G, Tripathi A, BenMarzouk-Hidalgo OJ, et al. Exploring *Anopheles* gut bacteria for *Plasmodium* blocking activity. Environ Microbiol. 2014;16:2980–2994.

Ramirez JL, Short SM, Bahia AC, Saraiva RG, Dong Y, Kang S, et al. *Chromobacterium Csp_P* reduces malaria and dengue infection in vector mosquitoes and has entomopathogenic and in vitro anti-pathogen activities. PLoS Pathog. 2014;10: e1004398.

Griffith BC, Weiss BL, Aksoy E, Mireji PO, Auma JE, Wamwiri FN, et al. Analysis of the gut-specific microbiome of field-captured tsetse flies, and its potential relevance to host trypanosome vector competence. 2018; Forthcoming.

Cirimotich CM, Ramirez JL, Dimopoulos G. Native microbiota shape insect vector competence for human pathogens. Cell Host Microbe 2011;10: 307–310.

Dennison NJ, Jupatanakul N, Dimopoulos G. The mosquito microbiota influences vector competence for human pathogens. Curr Opin Insect Sci. 2014;3: 6–13.

Fischer CN, Trautman EP, Crawford JM, Stabb EV, Handelsman J, Broderick NA. Metabolite exchange between microbiome members produces compounds that influence Drosophila behavior. Elife 2017;6. pii: e18855.

Podleśny M, Jarocki P, Wyrostek J, Czernecki T, Kucharska J, Nowak A, et al. *Enterobacter* sp. LU1 as a novel succinic acid producer - co-utilization of glycerol and lactose. Microb Biotechnol. 2017;10: 492–501.

Boissière A, Tchioffo MT, Bachar D, Abate L, Marie A, Nsango SE, et al. Midgut microbiota of the malaria mosquito vector *Anopheles gambiae* and interactions with *Plasmodium falciparum* infection. PLoS Pathog. 2012;8: e1002742.

Saraiva RG, Kang S, Simões ML, Angleró-Rodríguez YI, Dimopoulos G. Mosquito gut antiparasitic and antiviral immunity. Dev Comp Immunol. 2016;64: 53–64.

de Beer CJ, Moyaba P, Boikanyo SN, Majatladi D, Yamada H, Venter GJ, et al. Evaluation of radiation sensitivity and mating performance of *Glossina brevipalpis* males. PLoS Negl Trop Dis. 2017;11: e0005473.

McGraw EA, O’Neill SL. Beyond insecticides: new thinking on an ancient problem. Nat Rev Microbiol. 2013;11: 181–193.

Nurminsky DI, Nurminskaya MV, De Aguiar D, Hartl DL. Selective sweep of a newly evolved sperm-specific gene in *Drosophila*. Nature 1998;396: 572–575.

Nehme NT, Liégeois S, Kele B, Giammarinaro P, Pradel E, Hoffmann JA, et al. A model of bacterial intestinal infections in *Drosophila melanogaster*. PLoS Pathog. 2007;3: e173.

Dale C, Maudlin I. *Sodalis* gen. nov. and *Sodalis glossinidius* sp. nov., a microaerophilic secondary endosymbiont of the tsetse fly *Glossina morsitans morsitans*. Int J Syst Bacteriol. 1999;49 Pt 1: 267–275.

Vreysen MJ, Saleh KM, Ali MY, Abdulla AM, Zhu ZR, Juma KG, et al. *Glossina austeni* (Diptera: Glossinidae) eradicated on the island of Unguja, Zanzibar, using the sterile insect technique. J Econ Entomol. 2000;93: 123–135.

Vreysen MJ, Seck MT, Sall B, Bouyer J. Tsetse flies: their biology and control using area-wide integrated pest management approaches. J Invertebr Pathol. 2013;112 Suppl: S15–25.

Aksoy S, Weiss BL, Attardo GM. Trypanosome transmission dynamics in tsetse. Curr Opin Insect Sci. 2014;3: 43–49.

Song X, Wang M, Dong L, Zhu H, Wang J. PGRP-LD mediates *A. stephensi* vector competency by regulating homeostasis of microbiota-induced peritrophic matrix synthesis. PLoS Pathog. 2018;14: e1006899.

Baxter RH, Contet A, Krueger K. Arthropod innate immune systems and vector-borne diseases. Biochemistry 2017;56: 907–918.

Weiss B, Aksoy S. Microbiome influences on insect host vector competence. Trends Parasitol. 2011;27: 514–522.

Cirimotich CM, Dong Y, Clayton AM, Sandiford SL, Souza-Neto JA, Mulenga M, Dimopoulos G. Natural microbe-mediated refractoriness to *Plasmodium* infection in *Anopheles gambiae*. Science 2011;332: 855–858.

Dennison NJ, Saraiva RG, Cirimotich CM, Mlambo G, Mongodin EF, Dimopoulos G. Functional genomic analyses of *Enterobacter, Anopheles* and *Plasmodium* reciprocal interactions that impact vector competence. Malar J. 2016;15: 425.

Holmes P. Tsetse-transmitted trypanosomes--their biology, disease impact and control. J Invertebr Pathol. 2013;112 Suppl: S11–14.

Wang J, Weiss BL, Aksoy S. Tsetse fly microbiota: form and function. Front Cell Infect Microbiol. 2013;3: 69.

Aksoy E, Telleria EL, Echodu R, Wu Y, Okedi LM, Weiss BL, et al. Analysis of multiple tsetse fly populations in Uganda reveals limited diversity and species-specific gut microbiota. Appl Environ Microbiol. 2014;80: 4301–4312.

Geiger A, Fardeau ML, Njiokou F, Joseph M, Asonganyi T, Ollivier B, et al. Bacterial diversity associated with populations of *Glossina* spp. from Cameroon and distribution within the Campo sleeping sickness focus. Microb Ecol. 2011;62: 632–643.

Lindh JM, Lehane MJ. The tsetse fly *Glossina fuscipes fuscipes* (Diptera: Glossina) harbours a surprising diversity of bacteria other than symbionts. Antonie Van Leeuwenhoek 2011;99: 711–720.

Wang J, Wu Y, Yang G, Aksoy S. Interactions between mutualist *Wigglesworthia* and tsetse peptidoglycan recognition protein (PGRP-LB) influence trypanosome transmission. Proc Natl Acad Sci USA. 2009;106: 12133–12138.

Weiss BL, Wang J, Maltz MA, Wu Y, Aksoy S. Trypanosome infection establishment in the tsetse fly gut is influenced by microbiome-regulated host immune barriers. PLoS Pathog. 2013;9: e1003318.

Benoit JB, Attardo GM, Baumann AA, Michalkova V, Aksoy S. Adenotrophic viviparity in tsetse flies: potential for population control and as an insect model for lactation. Annu Rev Entomol. 2015;60: 351–371.

Wang J, Aksoy S. PGRP-LB is a maternally transmitted immune milk protein that influences symbiosis and parasitism in tsetse’s offspring. Proc Natl Acad Sci USA. 2012;109: 10552–10557.

MacLeod ET, Maudlin I, Darby AC, Welburn SC. Antioxidants promote establishment of trypanosome infections in tsetse. Parasitology 2007;134:827–831.

Vigneron A, Aksoy E, Weiss BL, Bing X, Zhao X, Awuoche EO, et al. A fine-tuned vector-parasite dialogue in tsetse’s cardia determines peritrophic matrix integrity and trypanosome transmission success. PLoS Pathog. 2018;14: e1006972.

Ridgley EL, Xiong ZH, Ruben L. Reactive oxygen species activate a Ca2+-dependent cell death pathway in the unicellular organism *Trypanosoma brucei brucei*. Biochem J. 1999;340: 33–40.

Neal-McKinney JM, Lu X, Duong T, Larson CL, Call DR, Shah DH, et al. Production of organic acids by probiotic lactobacilli can be used to reduce pathogen load in poultry. PLoS One 2012;7: e43928.

Buffie CG, Pamer EG. Microbiota-mediated colonization resistance against intestinal pathogens. Nat Rev Immunol. 2013;13: 790–801.

Nolan DP, Voorheis HP. Hydrogen ion gradients across the mitochondrial, endosomal and plasma membranes in bloodstream forms of *Trypanosoma brucei* solving the three-compartment problem. Eur J Biochem. 2000;267: 4601–4614.

Zilberstein D, Shapira M. The role of pH and temperature in the development of *Leishmania* parasites. Annu Rev Microbiol. 1994;48: 449–470.

Maltz MA, Weiss BL, O’Neill M, Wu Y, Aksoy S. OmpA-mediated biofilm formation is essential for the commensal bacterium *Sodalis glossinidius* to colonize the tsetse fly gut. Appl Environ Microbiol. 2012;78: 7760–7768.

Vigneron A, Weiss BL. Role of the microbiota during development of the arthropod vector immune system. In: Wikel S, editor. Arthropod vector: controller of disease transmission, vol. 1: vector microbiome and innate immunity of arthropods. Elsevier; 2017. pp. 161–169.

Michalkova V, Benoit JB, Weiss BL, Attardo GM, Aksoy S. Obligate symbiont-generated vitamin B6 is critical to maintain proline homeostasis and fecundity in tsetse flies. Appl Environ Microbiol. 2014;80: 5844–5853.

Aksoy E, Vigneron A, Bing X, Zhao X, O’Neill M, Wu YN, et al. Mammalian African trypanosome VSG coat enhances tsetse’s vector competence. Proc Natl Acad Sci USA. 2016;113:6961–6966.

Weiss BL, Savage AF, Griffith BC, Wu Y, Aksoy S. The peritrophic matrix mediates differential infection outcomes in the tsetse fly gut following challenge with commensal, pathogenic, and parasitic microbes. J Immunol. 2014;193:773–782.

Weiss BL, Maltz M, Aksoy S. Obligate symbionts activate immune system development in the tsetse fly. J Immunol. 2012;188: 3395–3403.

Narasimhan S, Fikrig E. Tick microbiome: the force within. Trends Parasitol. 2015;31: 315–323.

Welburn SC, Arnold K, Maudlin I, Gooday GW. Rickettsia-like organisms and chitinase production in relation to transmission of trypanosomes by tsetse flies. Parasitology 1993;107:141–145.

